# Apo and ligand-bound high resolution Cryo-EM structures of the human Kv3.1 reveal a novel binding site for positive modulators

**DOI:** 10.1101/2021.07.13.452180

**Authors:** Mathieu Botte, Sophie Huber, Denis Bucher, Julie K. Klint, David Rodríguez, Lena Tagmose, Mohamed Chami, Robert Cheng, Michael Hennig, Wassim Abdul Rahman

## Abstract

Kv3 ion-channels constitute a class of functionally distinct voltage gated ion channels characterized by their ability to fire at a high frequency. Several disease relevant mutants, together with biological data, suggest the importance of this class of ion channels as drug targets for CNS disorders, and several drug discovery efforts have been reported. Despite the increasing interest for this class of ion channels, no structure of a Kv3 channel has been reported yet. We have determined the cryo-EM structure of Kv3.1 at 2.6 Å resolution using full length wild type protein. When compared to known structures for potassium channels from other classes, a novel domain organization is observed with the cytoplasmic T1 domain, containing a well resolved Zinc site and displaying a rotation by 35°. This suggests a distinct cytoplasmic regulation mechanism for the Kv3.1 channel. A high resolution structure was obtained for Kv3.1 in complex with a novel positive modulator Lu AG00563. The structure reveals a novel ligand binding site for the Kv class of ion channels located between the voltage sensory domain and the channel pore, a region which constitutes a hotspot for disease causing mutations. The discovery of a novel binding site for a positive modulator of a voltage gated potassium channel could shed light on the mechanism of action for these small molecule potentiators. This finding could enable structure-based drug design on these targets with high therapeutic potential for the treatment of multiple CNS disorders.

## I. Introduction

The characteristic electrical activity of neurons and their ability to conduct, transmit, and receive electric signals, results from the opening and closing of ion channels in the neuron plasma membrane.

Voltage-gated potassium channels (Kv) form a family of transmembrane ion channels which are voltage-sensitive and selective for potassium ions. The Kv3 family consists of Kv3.1 (*KCNC1*), Kv3.2 (*KCNC2*), Kv3.3 (*KCNC3*) and Kv3.4 (*KCNC4*). Kv3.1 channels conduct a delayed rectifying outward current and have a high activation threshold (≈ -20 mV) as well as rapid activation and deactivation kinetics (Kaczmarek and Zhang, 2017). The Kv3.1 channels are thereby responsible for neuronal repolarization, and high-frequencey action potential firing (Rudy and McBain, 2001; Wang et al., 1998).

The Kv3 channels, like other Kv channels, are tetrameric structures of four pore-forming alpha subunits. Each alpha subunit is comprised of six transmembrane helixes (S1-6). The S4 helix has positively charged residues at every third residue, and acts as a voltage sensor that together with S1-3 is denoted the voltage sensing domain (VSD, S1-4). The sensing of voltage triggers a conformational change that opens the channel pore domain (PD, S5-6) and allows a flux of K^+^ ions through the channel.

Kv channels often associate with accessory beta subunits like Kvβ1, 2 and 3 (Rettig et al., 1994; Morales et al., 1995; Majumder et al., 1995; England et al., 1995). In some cases these subunits are involved in N-type inactivation of the channel, as for Kv1.2 regulation by Kvβ1 (Rettig et al., 1994). It was shown that neither Kvβ1 nor Kvβ2 co-immuno-precipitates with Kv3.1 (K. Nakahira, G. Shi, K. Rhodes, 1996) pointing to a distinct regulatory mechanism of the Kv3 family of ion-channels. Other potential accessory subunits such as Mink, Mirp1 and Mirp2 were shown to modulate the function of Kv3.1 and Kv3.2 by slowing their activation (Lewis et al., 2004). In the case of Mirp2 it was suggested that it co-assembles with Kv3.1 in the rat brain, but not in a universal manner. For example no association between Kv3.1 and Mirp2 could be observed in rat E18 hippocampal neurons (McCrossan et al., 2003). Overall, there is no clear report of Kv3.1 regulation by accessory subunits.

Importance of the Kv3 subclass of potassium channels for CNS related diseases is supported by identification of several disease mutations connected to rare forms of epilepsy (Oliver et al., 2017; Muona et al., 2015). In addition to the high therapeutic potential in seizure treatment, Kv3 modulators might rescue dysfunction and alteration in gamma oscillations. These modulators could regulate diverse cognitive functions such as sensory integration, attention and working memory (Andrade-Talavera et al., 2020), which are for example attenuated in Alzheimer’s disease (Uhlhass et al., 2006). Consequently, several Kv3 small molecule modulators with different profiles and properties have been identified. AUT1 (Rosato-Siri et al., 2015) and AUT2 (Brown et al., 2016) have been reported as positive modulators of Kv3.1 and Kv3.2 channels. In addition, H. Lundbeck A/S has discovered a series of Kv3.1 modulators (Sams et al., 2020) that act as potentiators by lowering the voltage threshold for activation, leading to an increased peak current.

Here, we report the first cryo-EM structures of a Kv3 channel in apo form and in complex with the Lu AG00563 potentiator ligand. Analysis of the structures give insights into the tetramer association and identifies a novel potentiator binding site of the Kv3.1 channel.

## II. Results

### a. Protein biochemistry and apo protein EM structure

Construct design was done to ensure optimal relevance of the protein for structure-based drug design. In the case of Kv3.1, two splice isoforms are known, the canonical Kv3.1a isoform and Kv3.1b isoform which has an extension at the C-terminal (Luneau et al., 1991). Both isoforms display identical basal currents but Kv3.1b function is further regulated by a phosphorylation site at the C-terminal (Macica et al., 2003). Consequently, the selection of the isoform is not critical for structure-based drug design targeting the core channel. Expression in HEK293 and purification were performed with the canonical full-length wild type Kv3.1a isoform tetramer referred to as flWT-Kv3.1. Biochemical analysis of the alpha subunit showed no co-purification with any endogenous subunit at a level which could be detected by Coomassie staining. High level of homogeneity and detergent stability were achieved as judged by the size exclusion profile and negative staining analysis of the purified sample (SupFig 1).

The cryo-EM structure of flWT-Kv3.1 at 2.65 Å overall resolution (SupFig 2) enabled *de novo* model building (SupFig 3). Human flWT-Kv3.1 adopts a tetrameric swapped organization, with the VSD of each subunit interacting with the PD of the neighboring subunit (Fig 1).

**Figure 1.**
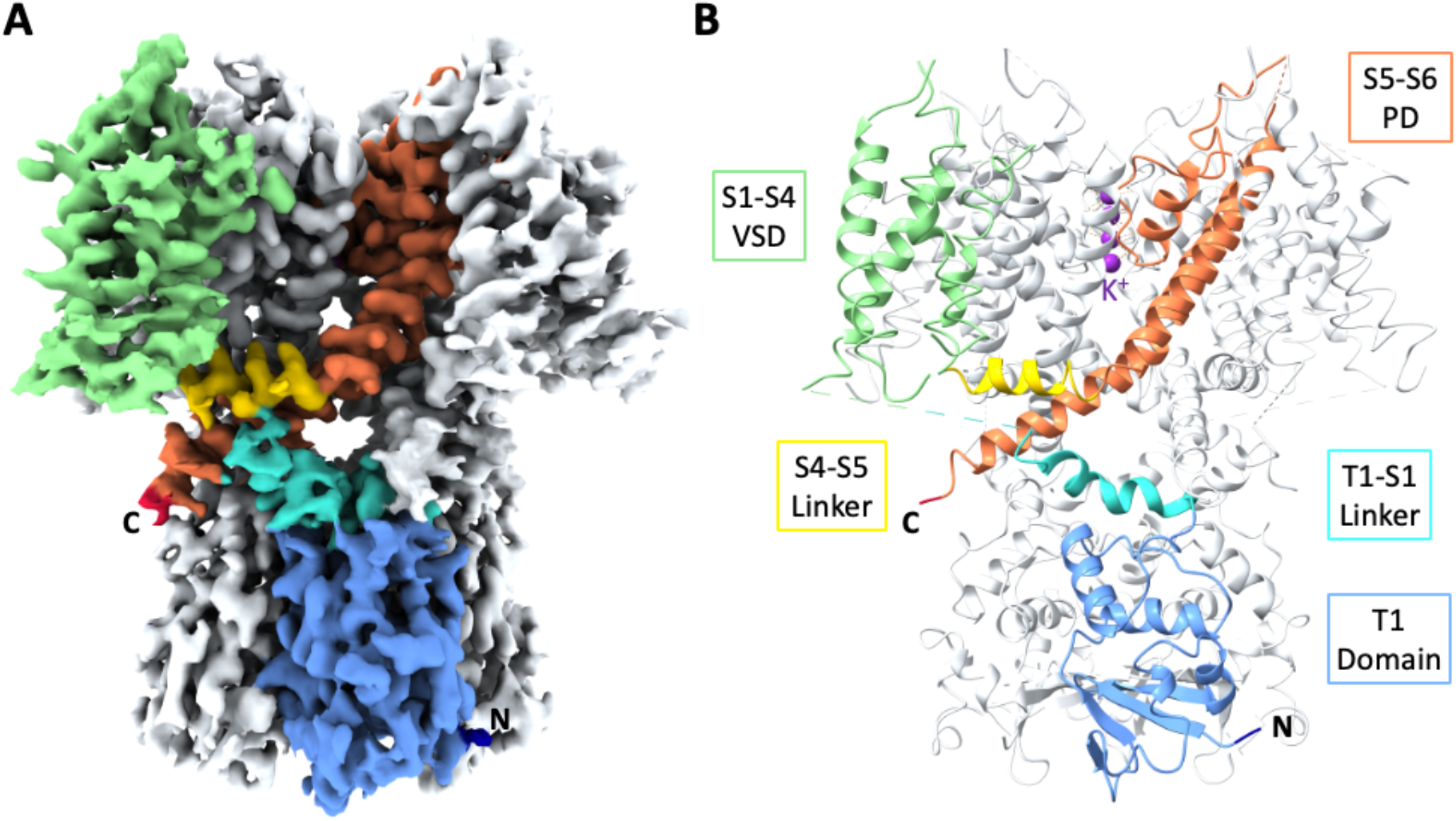
Structure of apo flWT-Kv3.1. **A**. Cryo-EM reconstruction of apo flWT-Kv3.1. **B**. Atomic model of apo flWT-Kv3.1. The color code is identical in both figures and highlights the different domains and their organization in flWT-Kv3.1. For clarity, only one monomer has been colored, the remaining three monomers are colored in light grey. The N-termini end is colored in dark blue, the T1 domain in light blue, the T1-S1 linker in cyan, the voltage sensing domain (VSD) in light green, the S4-S5 linker in yellow, the pore domain (PD) in orange and the C-termini end in red. Potassium ions visible in the reconstruction were modeled and are colored in purple.

Overall, the transmembrane domain organization is similar to what was observed with Kv channels in previously published structures, particularly for Kv1.2 (Long et al., 2007; Long et al., 2005a; Long et al., 2005b; Matthies et al., 2018) which shares the highest sequence similarity with Kv3.1. In contrast, the T1 cytoplasmic domain shows a novel structural feature by adopting a 35° twisted conformation (Fig 2A,B). Furthermore, the T1 domain tetramerization of Kv3.1 is mediated by a Zn^2+^ atom coordinating interaction between His77, Cys104 and Cys105 from one polypeptide and Cys83 from the neighboring polypeptide (Fig 2C) similar to the structure of the T1 domain tetramer of Kv4.2 (Scannevin et al., 2004).

**Figure 2.**
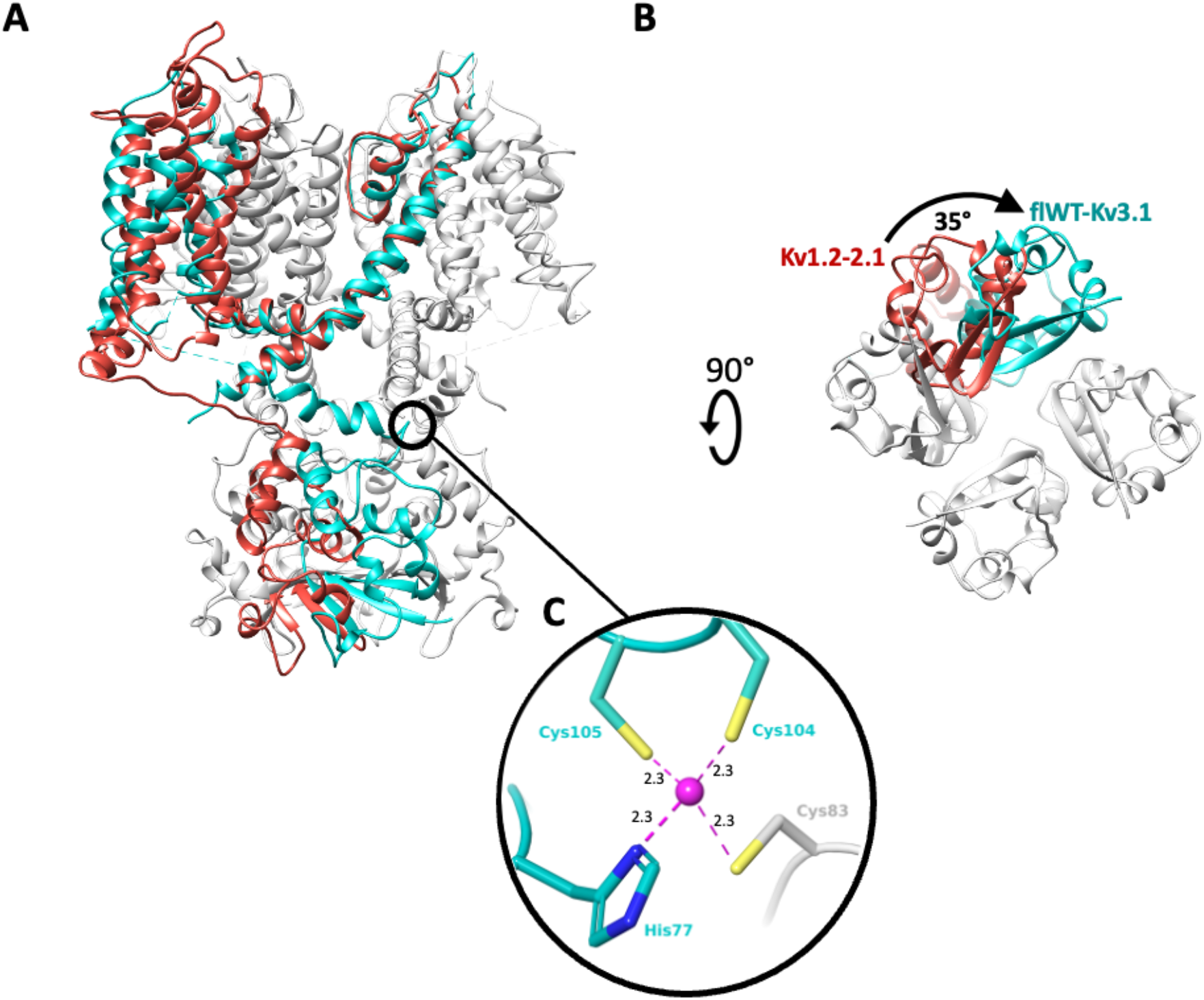
T1 cytoplasmic domain adopting a 35° twisted conformation. **A**. Overlay of a monomer of Kv1.2-2.1 paddle chimera structure (red, pdb code: 6ebk) with a tetramer of apo flWT-Kv3.1 (cyan, only one monomer colored). The overlay is done on the pore domain. **B**. apo flWT-Kv3.1 and Kv1.2-2.1 structures are shown from an intracellular viewpoint to illustrate the 35° twisting motion. **C**. Coordination site for Zn^2+^ atom (purple, and distances to nearest heavy atom reported in Angstroms) in flWT-Kv3.1. The Zn-site appears to assist the T1 tetramerization by bridging His77 (T1), Cys104 (T1-S1 linker) and Cys105 (T1-S1 linker) from one monomer, with Cys83 (T1) from a neighboring monomer.

As observed with other Kv channels, EM density is visible in the center of the selectivity filter (Fig 1B) corresponding to the average density of distinct K^+^ ions. Four sites of potassium ions could be modeled coordinating the residues forming the selectivity filter. Before entering the Kv family-conserved selectivity filter, K^+^ ions pass through the lower gate. The analysis of the pore radius in this region shows that the lower gate of the channel is in an open conformation with a diameter of 3.3 Å (SupFig4), similar to other structures solved in the open state, such as Kv1.2 (Long et al., 2007; Long et al., 2005a; Long et al., 2005b; Matthies et al., 2018) and Kv7.1 (Sun and MacKinnon, 2017; Sun and MacKinnon, 2020) with similar pore radius at the most constricted region.

### b. EM Structure with potentiator ligand

H. Lundbeck A/S identified a series of novel compounds which act as Kv3 channel potentiators by shifting the activation threshold to the hyperpolarized direction (Sams et al., 2020). The measured ECΔ5mV parameter corresponds to the effective concentration needed to shift the activation threshold by 5mV towards the hyperpolarized direction. Within the series, Lu AG00563 (Ex86), for which the measured ECΔ5mV is 3 µM, showed a good aqueous solubility in our buffer system and could be added to the purified protein at a final concentration of 500 µM without impairing the quality of the sample preparation. The EM structure with the ligand bound was determined at an overall resolution of 3.0Å (SupFig 5). The organization of the domains and subunits and all the observations made for the apo structure are identical. No variation of the pore radius could be observed and the lower gate does not display any structural rearrangement. After careful analysis of the cryo-EM reconstruction for non-protein densities, the density previously observed for the K^+^ ions in the apo-structure were confirmed as well. Furthermore, an additional density, absent in the apo-structure, was observed (Fig. 3A). This density is located at the interface between two monomers where the VSD of one monomer, is interacting with the PD of the neighboring subunit (Fig. 3A,B). Lu AG00563 could be fitted into the density in an unambiguous conformation (Fig. 3B), either manually or assisted by computational docking methods. Analysis of the ligand binding cavity highlights interactions that could be explored for the design of modulators of Kv3 channels (Fig. 3C,D). A favorable geometry for a hydrogen bond interaction is observed between the side chain of Thr325, located next to S4 at the VSD-PD interface, and the ligand carbonyl group. Several hydrophobic residues appear to contribute to the ligand interaction. In particular residues from the S1 helix like Ala193, Phe194, Leu197, Leu201, as well as Leu324 and Phe328 of helix S4, and residues Ile350 and Leu354 of helix S5 from the neighboring monomer.

**Figure 3.**
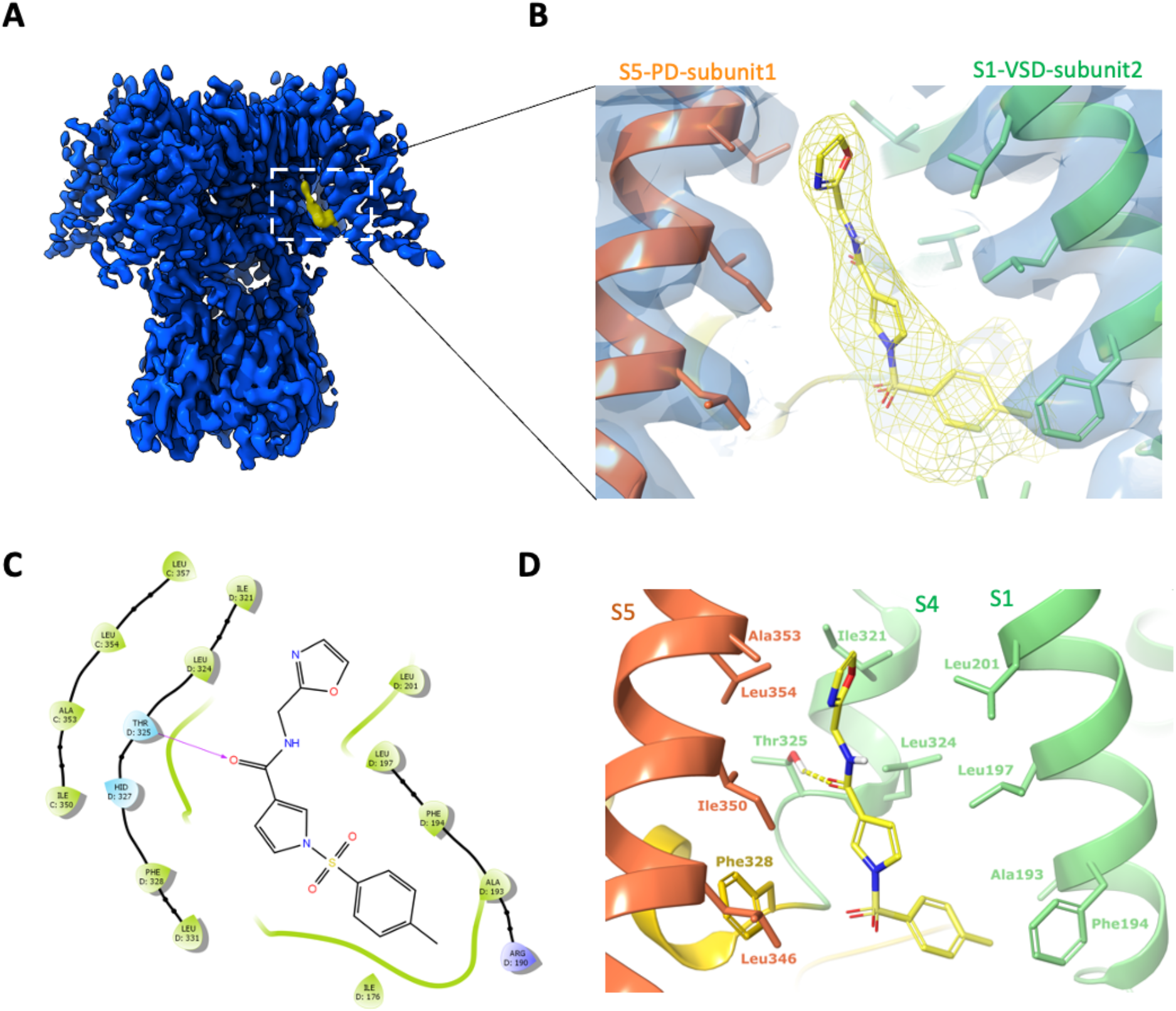
Binding site observed for the Lu AG00563 positive modulator. **A**. Cryo-EM density map (blue) with region of non protein density attributed to the ligand (yellow). **B**. Close-up view of the ligand binding pocket, displaying both the ligand and protein experimental densities. **C**. Simplified diagram of the protein-ligand interactions, highlighting the H-bond with Thr325, hydrophobic residues (green) and polar residues (cyan). **D**. Tri-dimensional representation of interactions in the binding site. The S1 and S4 helices are shown in light green, S5 in orange, and the S4-S5 linker in yellow.

## III. Discussion and conclusion

### Structural differences of Kv3.1 in comparison to other Kv channels

Among the Kv channels for which a structure is available Kv3.1 shares the highest sequence and structure similarity with Kv1.2 (Long et al., 2007; Long et al., 2005a; Long et al., 2005b; Matthies et al., 2018). The major structural difference between Kv1.2 and Kv3.1 is observed at the intracellular T1 domain with a 35° twist between the two T1 domains of both channels. Another major dissimilarity is the zinc binding site at the base of the T1 tetramerization which is not observed in Kv1.2. These structural differences between the Kv1.2 and Kv3.1 channels seem to be mainly driven by different regulation mechanisms occurring at the intracellular side. Kv1.2 is regulated at the intracellular side by its association via the T1 domain to Kvβ1 which is involved in N-type inactivation of the channel. The lack of confirmation of any association of Kv3.1 with Kvβ1 suggests that the activity regulation of the channel is performed with distinct mechanisms. The structural data suggest for Kv3.1 channels a mechanism with zinc mediated T1 tetramerization of the cytoplasmic regulation that may be controlled by Zinc concentration.

### A novel Kv potentiator-binding site

The binding site for Lu AG00563 at the interface between the VSD and the PD represents a key finding of the present structure. Lu AG00563 displays similar electrophysiological behavior to retigabine, which is a potentiator inducing a negative shift in the required activation voltage. The structure of Retigabine bound to Kv7.2 (Li et al., 2021b) and Kv7.4 (Li et al., 2021a) was recently solved and shows that the binding site of the compound is located next to the PD between S5 from one subunit and S6 from the neighboring subunit (Fig. 4B). In contrast, Lu AG00563 binds to Kv3.1 between the VSD of one subunit, and the PD of the neighboring subunit (Fig3 and Fig4A). Therefore, Lu AG00563 and retigabine display similar functional properties by binding to two distinct pockets. Hence, the current structures offer a new insight for how these channels could be modulated by small molecules.

**Figure 4.**
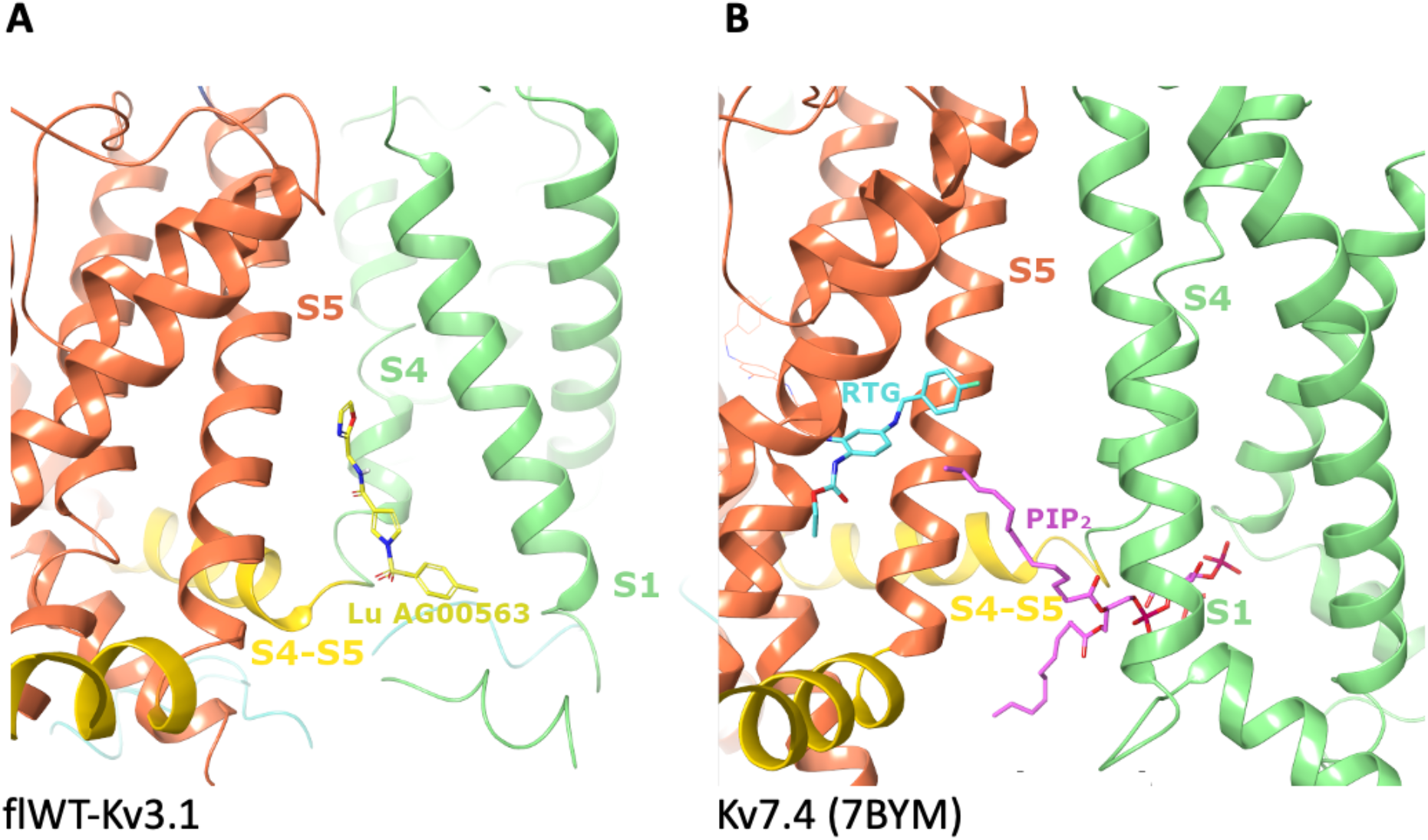
Comparison of the binding sites for Lu AG00563, in flWT-Kv3.1, and PIP*2* and retigabine, in Kv7.4. **A**. Binding site discovered for Lu AG00563 at the interface between S1, S4 and the S4-S5 linker. The S1 and S4 helices are shown in light green, S5 in orange, and the S4-S5 linker in yellow. Lu AG00563 is shown in yellow. **B**.Comparison with the Kv7.4 structure (pdb code: 7bym), highlighting similarities with the PIP2 binding site showing interactions with S1, S4 and the S4-S5 linker. In contrast, the retigabine (RTG) binding site is located next to the pore (S5) domain. PIP2 is shown in purple and retigabine is shown in Cyan.

The newly discovered site for Lu AG00563 is located in close proximity to the S4 segment, which is paved with positively charged residues acting as voltage sensors. However, the ligand is not fully entering the S1-S4 bundle. Interestingly, the structure of Kv7.4 bound to PIP2 carrying a lipid headgroup entering the S1-S4 bundle (Fig 4B) shows a related binding site for this endogenous lipid, and has been reported to favor the opening of Kv7 channels (Rodriguez-Menchaca et al., 2012). The binding site is also a a hotspot for several disease related mutations that were reported in the Kv3 class, highlighting the functional importance of the binding site. Some of the mutants occur in the S4 segment, which is strictly conserved within the Kv3 subclass (SupFig 6a). In particular, point mutations occurring in the VSD that have been found in patients with spinocerebellar ataxia are associated with current amplitude reduction. Some of these mutations involve positive-charge neutralization occurring in S4, such as Kv3.3-R420H (Gallego-Iradi et al., 2014) or Kv3.1-R320H and Kv3.3-R423H (Park et al., 2019). Of particular interest is the T428I mutation observed in Kv3.3 (Németh et al., 2013), since Thr428 is structurally equivalent to Thr325 in Kv3.1 which forms the key hydrogen bond interaction with Lu AG00563 (SupFig 6B). Taken together, these observations support a role for residues near the Lu AG00563 binding site area in the modulation of the channel activity (SupFig 6B).

### Impact to drug discovery

The binding site of the Lu AG00563 potentiator at the bottom of S4 and next to the S4-S5 linker has only been previously observed for the non-drug like lipid PIP2 in Kv7.4. The hydrogen bond of the carbonyl of Lu AG00563 with the hydroxyl of Thr325 appears to be a key interaction for this binding mode, as it uses the only polar side chain residue present in this inter-subunit area between helices S4 and S5. The lack of high-resolution structures for Kv3 channels has previously hindered the development of potentiators by structure-based methods. In particular, Kv3.1 is known as a potential drug target for the treatment of multiple CNS related disorders. The ion channel structure reported here could open up new opportunities for the design of drug molecules with enhanced properties and offers an excellent starting point to study the druggability of alternative pockets for the discovery and characterization of Kv3.1 modulators.

## Methods

### Expression of Kv3.1 in HEK293 F cells

The cDNA of the wild type full length human Kv3.1 isoform a (flWT-Kv3.1 with uniprot reference P48547) with a Carboxy-terminal tag composed of prescission 3C cleavage site followed by GFP was cloned in the expression plasmid pLXBM7 which allows expression of the target protein in mammalian cells with the control of the CMV promoter. The plasmid was produced in large amounts by using the NucleoBond PC 10000 EF kit. HEK293F cells were transfected using PEI 25K linear transfection reagent (Polysciences). Expression was performed at 37°C for 7 hours followed by a second expression phase at 30°C for 72 hours.

### Purification of flWT-Kv3.1 tetramer

To purify the flWT-Kv3.1 tetramer, cells were resuspended in buffer A (Tris 20 mM pH 7.5, KCl 200 mM, MgCl2 1 mM, Lauryl Maltose Neopentyl Glycol 1%, cholesteryl hemisuccinate 0.1 %, Protease inhibitor cocktail EDTA free from Roche according to manufacturer recommendations and DNASE from Roche at 0.01 mg/ml). Cells were homogenized and the flWT-Kv3.1 tetramer was extracted by gentle stirring for 2 hours at 4°C. The non soluble fraction was removed by ultra-centrifugation at 42000 rpm at 4°C for 1 hour using Ti45 Beckman rotor. The soluble fraction was mixed with non commercial Pro-GFP resin (anti-GFP binder coupled to CNBr Activated Sepharose resin from GE-healthcare) pre-equilibrated with buffer B (Tris 20 mM pH 7.5, KCl 200 mM, Lauryl Maltose Neopentyl Glycol 0.01% and DTT 1mM). After incubation for 2 hours at 4°C the resin was washed extensively with buffer B and the flWT-Kv3.1 tetramer was eluted by addition of 100 µl of HRV 3C Protease at 1U/µl concentration from Takara Bio. Elution was performed overnight at 4°C. Eluted protein was concentrated on vivaspin 100 kDa cutoff Vivaspin Turbo 15 concentrator and injected on a 10/30 Superose 6 size exclusion chromatography column pre-equilibrated with buffer B. Fractions which correspond to the tetramer were pooled and concentrated to reach 1mg/ml concentration.

### Computational Methods

All modelling was done in Maestro (Schrodinger inc.). Docking of Lu AG00563 was performed with GlideEM (Schrodinger Maestro version 2020-1) in its standard parametrization (Schrödinger Release 2020-1: Glide, Schrödinger, LLC, New York, NY, 2021) assuming a local EM map resolution of 3 Å. The initial sampling phase generated a large number of candidate poses that were then filtered into the five top scoring poses for the refinement phase and visual inspection. Real-space refinement was then performed using the software Phenix and the OPLS3e/VSGB2.1 force field. Following refinement, an unambiguous pose could be selected based on its docking score, the additional electron density explained by the ligand (density score), and visual inspection of chemical interactions between the ligand and the protein environment (Robertson et al., 2020, Adams et al., 2010, Roos et al., 2019).

### Sample preparation and EM analysis

Negative stain electron microscopy showed good size homogeneity, optimal particles distribution and shape of a typical potassium channel molecule. Subsequently, sample conditions were screened by cryo-EM for the optimal concentration, grids and freezing parameters. After evaluation of the conditions, quantifoil (2/2) 300-mesh copper grids were glow-discharged for 50 seconds prior sample freezing. 3µl of flWT-Kv3.1 in presence or not of 500µM Lu AG00563 at a concentration of 2mg/ml were placed on the grid, blotted for 3.0s and flash frozen in a mixture of liquid propane and liquid ethane cooled with liquid nitrogen using a Vitrobot Mak IV (FEI) operated at 10°C and 100% humidity.

The EM data collection statistics in this study is reported in table 1. Data were recorded on a FEI Titan Krios microscope operated at 300 kV equipped with a K2 Summit direct electron detector (Gatan Inc.) and Quantum-LS energy filter (slit width 20 eV ; Gatan Inc.). The automation of the data collection was done with the software SerialEM (Mastronarde, 2005). Movies were recorded in electron-counting mode fractionating 70 electrons over 40 frames or 60 electrons over 40 frames for, respectively, the flWT-Kv3.1 (apo) or the flWT-Kv3.1 with Lu AG00563 samples. A defocus range of -0.8 to -2.8 µm was used and the physical pixel size was 0.82 Å/pixel. All the movies were gain-normalized, motion-corrected and dose-weighted with MotionCor2 (Zheng et al., 2017). The micrographs were sorted using FOCUS (Biyani et al., 2017) to clear up those unwell images.

**Table 1:**
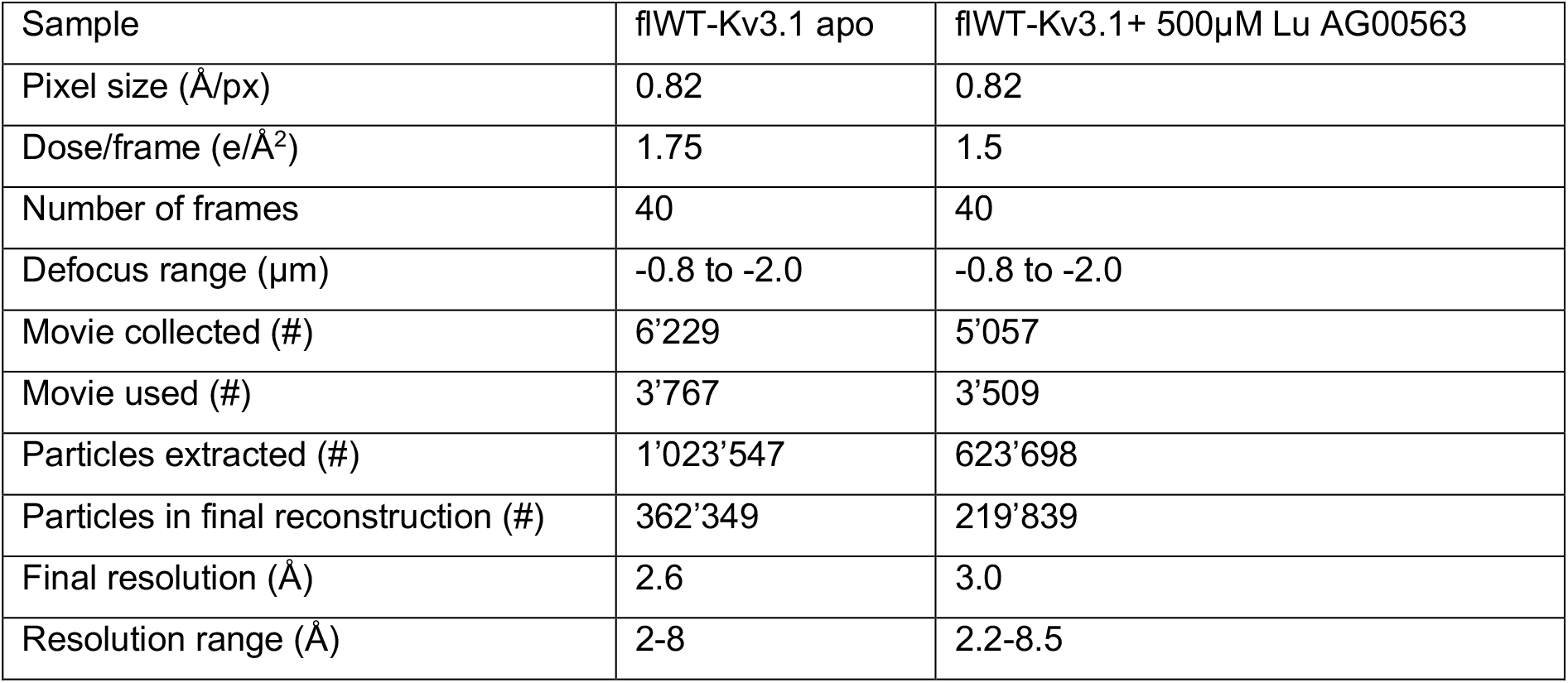
Data collection parameters and dataset statistics.

### Image processing

The following processing workflows were used for the samples in the study. The aligned movies were imported into CryoSPARC V2 (Punjani et al., 2017, Rohou & Grigorieff, 2015). A set of aligned averages with a calculated defocus range of −0.8 to –2.8 μm was selected from which averages with poor CTF estimation statistics were discarded. Automated particle picking in CryoSPARC V2 resulted in 1’023’547 and 623’698 particle locations for respectively, the apo sample and the sample in presence of Lu AG00563. After several rounds of 2D classification, particles were selected and subjected to 3D classification using the multi class ab initio refinement process and heterogenous refinement. The best resolved classes consisting of 362’349 particles for the apo sample, and 219’839 particles for the Lu AG00563 sample were finally subjected to 3D non-uniform refinement. The overall resolution of the resulted map was estimated at 2.65Å for apo flWT-Kv3.1 and 3.03Å for Lu AG00563-flWT-Kv3.1 based on the Fourier shell correlation (FSC) at 0.143 cutoff (Scheres & Chen, 2012).

### Model building and refinement

An initial flWT-Kv3.1 was generated using SWISS-MODEL using as templates the Kv1.2 structure (PDB-ID XXXX). Rigid body fitting was initially done in Chimera followed by manually rebuilding of the model in Coot. Remaining clashes between sidechains were detected using Schrodinger version 2019-4, and remodelled using prime (Zhu et al., 2014). Manual inspection of missing H-bonds in the model was used to refine sidechain positions. Finally, real-space refinement was performed in Phenix version 1.17-3644, applying Ramachandran plot restraints (Liebschner et al., 2019).

---

**SupFig 1.**
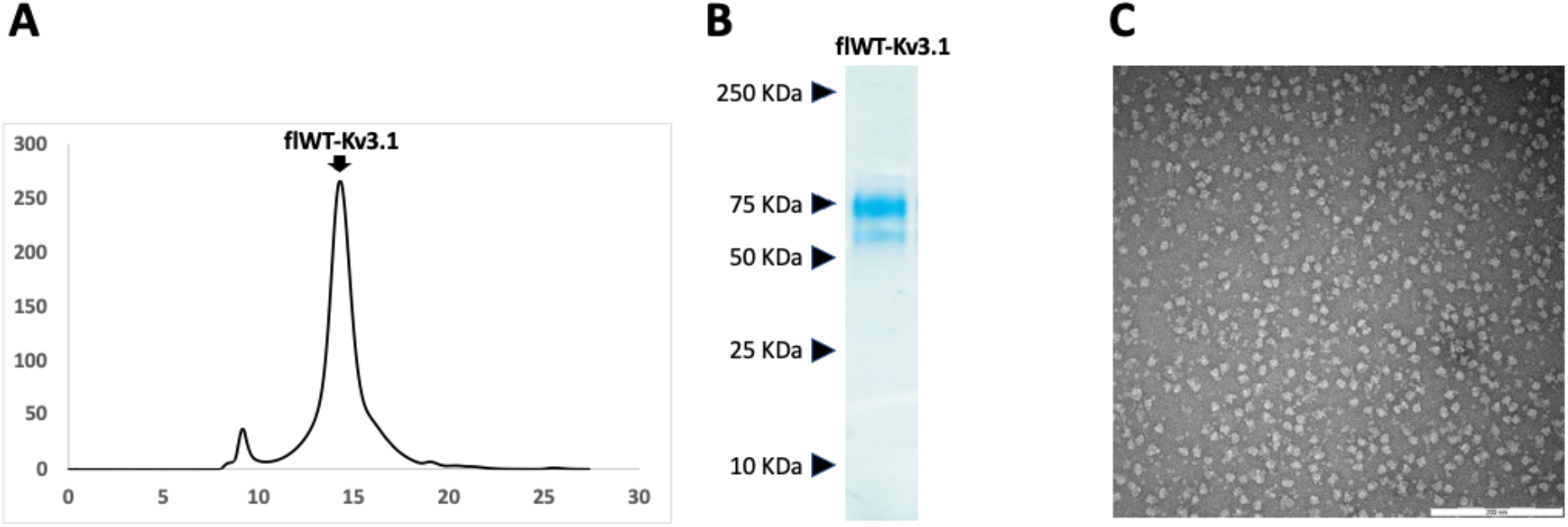
Purification and negative staining analysis of flWT-Kv3.1 tetramer. **A**. After affinity purification the protein sample was subjected to superose 6 Size Exclusion chromatography. The peak corresponding to the tetramer is indicated with an arrow. **B**. The final sample was analyzed by SDS-PAGE. Positions of the protein markers bands are indicated with arrows on the left side. **C**. The final protein sample was analyzed by negative staining.

**SupFig2.**
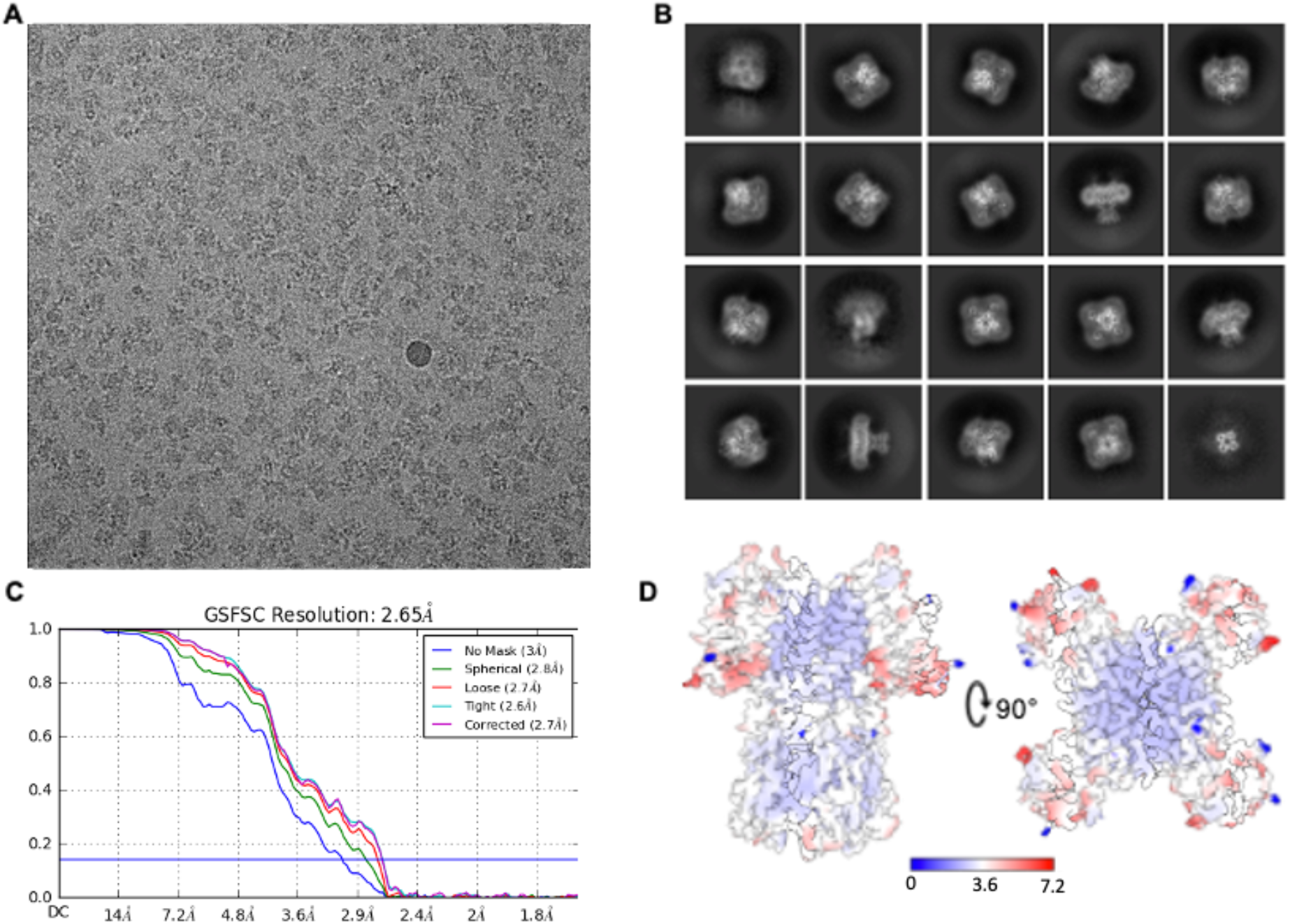
Cryo-EM acquisition and map of the apo flWT-Kv3.1. **A**. Example of a micrograph of vitrified apo flWT-Kv3.1. **B**. Example of 2D class averages. **C**. Gold standard Fourier-Shell correlation resolution plot with a cut-off at 0.143 indicated by a blue line. **D**. Map of the apo flWT-Kv3.1 with color-coded local resolution.

**SupFig3.**
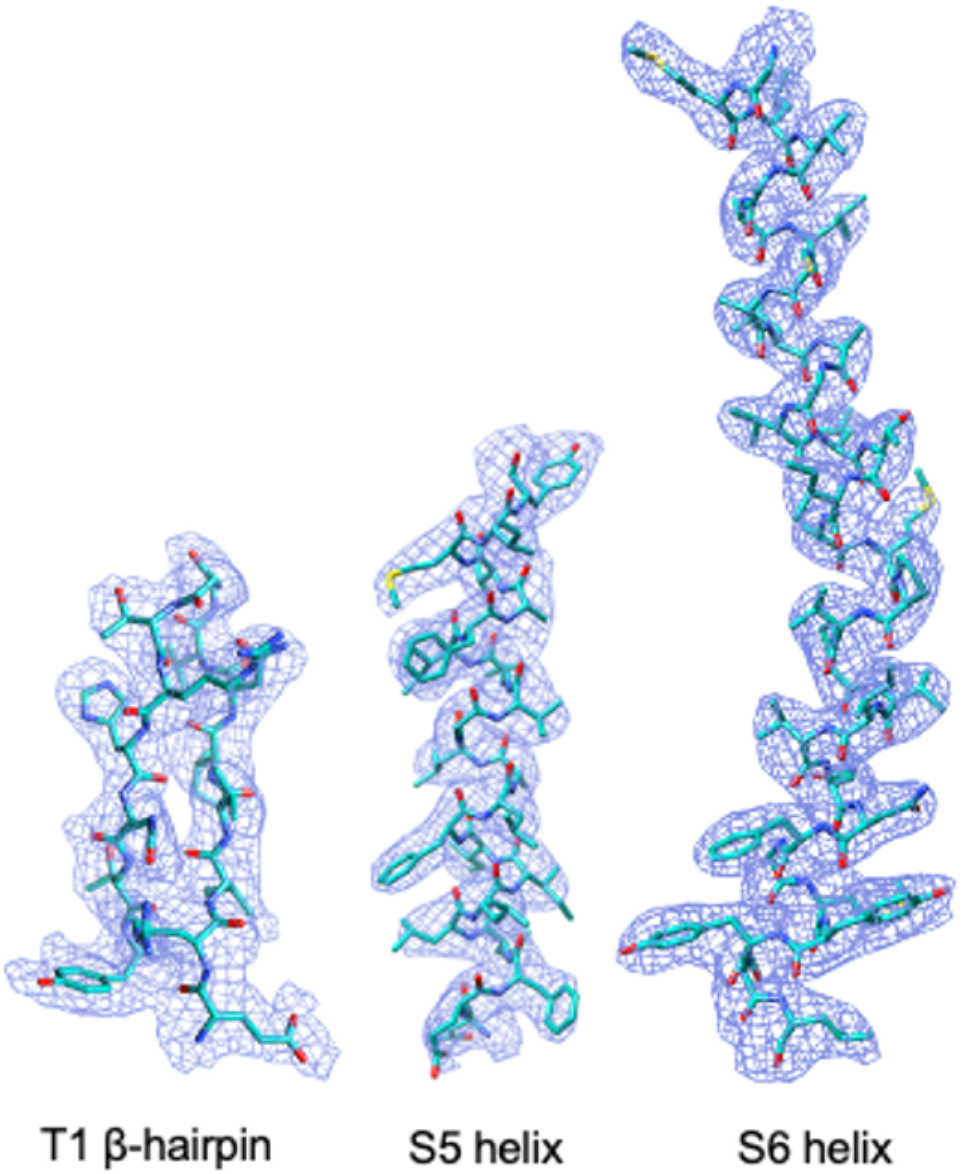
Agreement of the Cryo-EM map and the corresponding model. Comparison of the cryo-EM map quality versus the atomic model for different part of the protein. The cryo-EM map is represented as blue mesh and the atomic model is represented in cyan.

**SupFig4.**
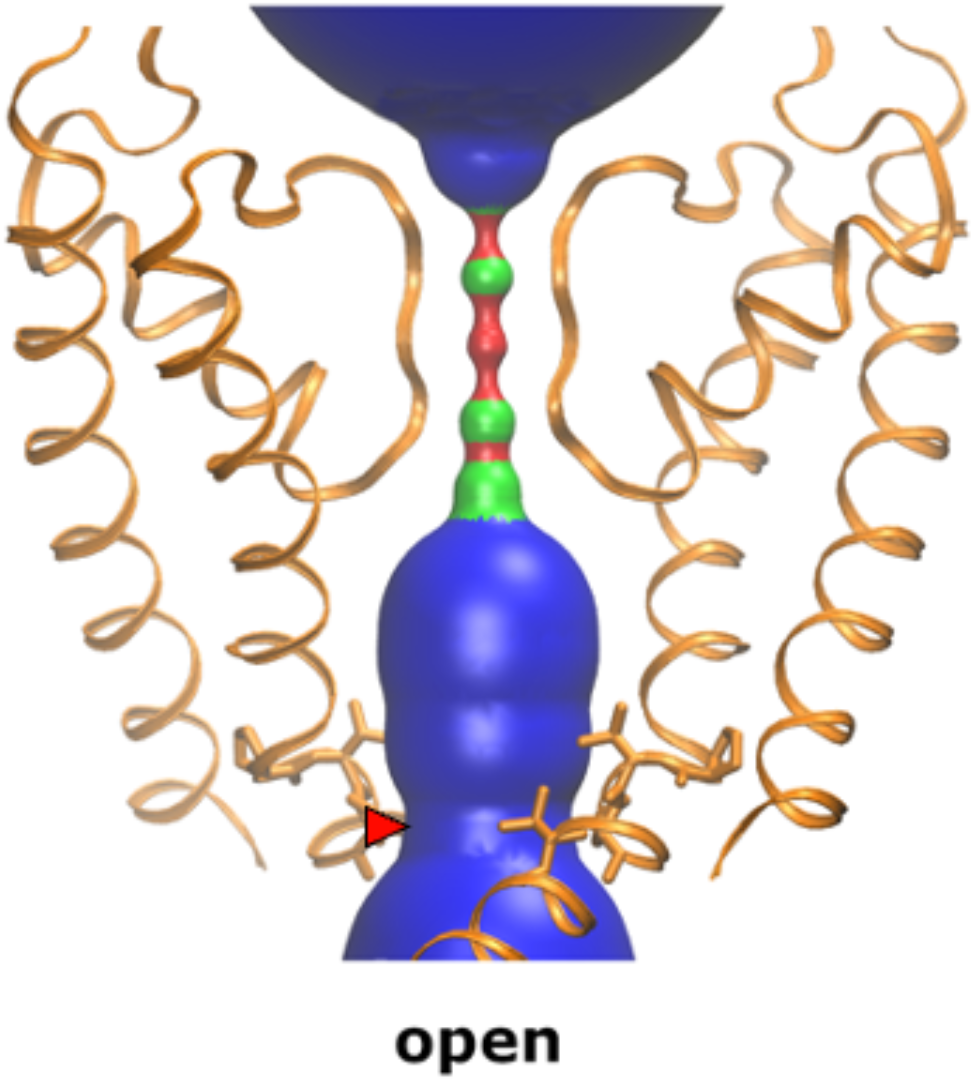
Cartoon representation of the cylindrical radius within the pore. The pore radius is shown as calculated with the HOLE2 program, indicating an open channel at the lower gate region indicated with a red arrow.

**SupFig5.**
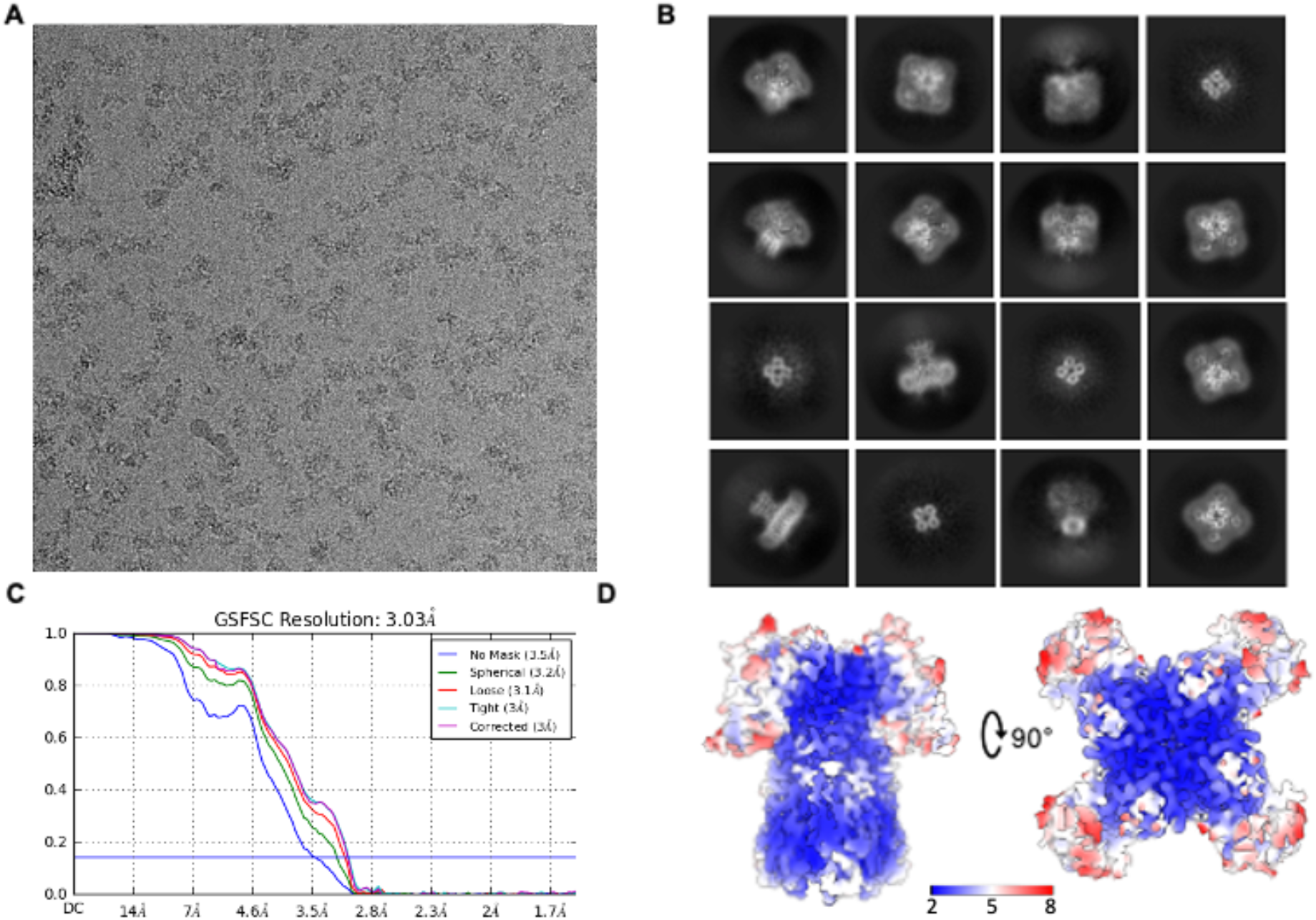
Cryo-EM acquisition and map of flWT-Kv3.1 in presence of Lu AG00563. **A**. Example micrograph of vitrified flWT-Kv3.1 in presence of Lu AG00563. **B**. Example of 2D class averages. **C**. Gold standard Fourier-Shell correlation resolution plot with a cut-off at 0.143 indicated by a blue line. **D**. Map of the flWT-Kv3.1 in presence of Lu AG00563 with color-coded local resolution.

**SupFig 6.**
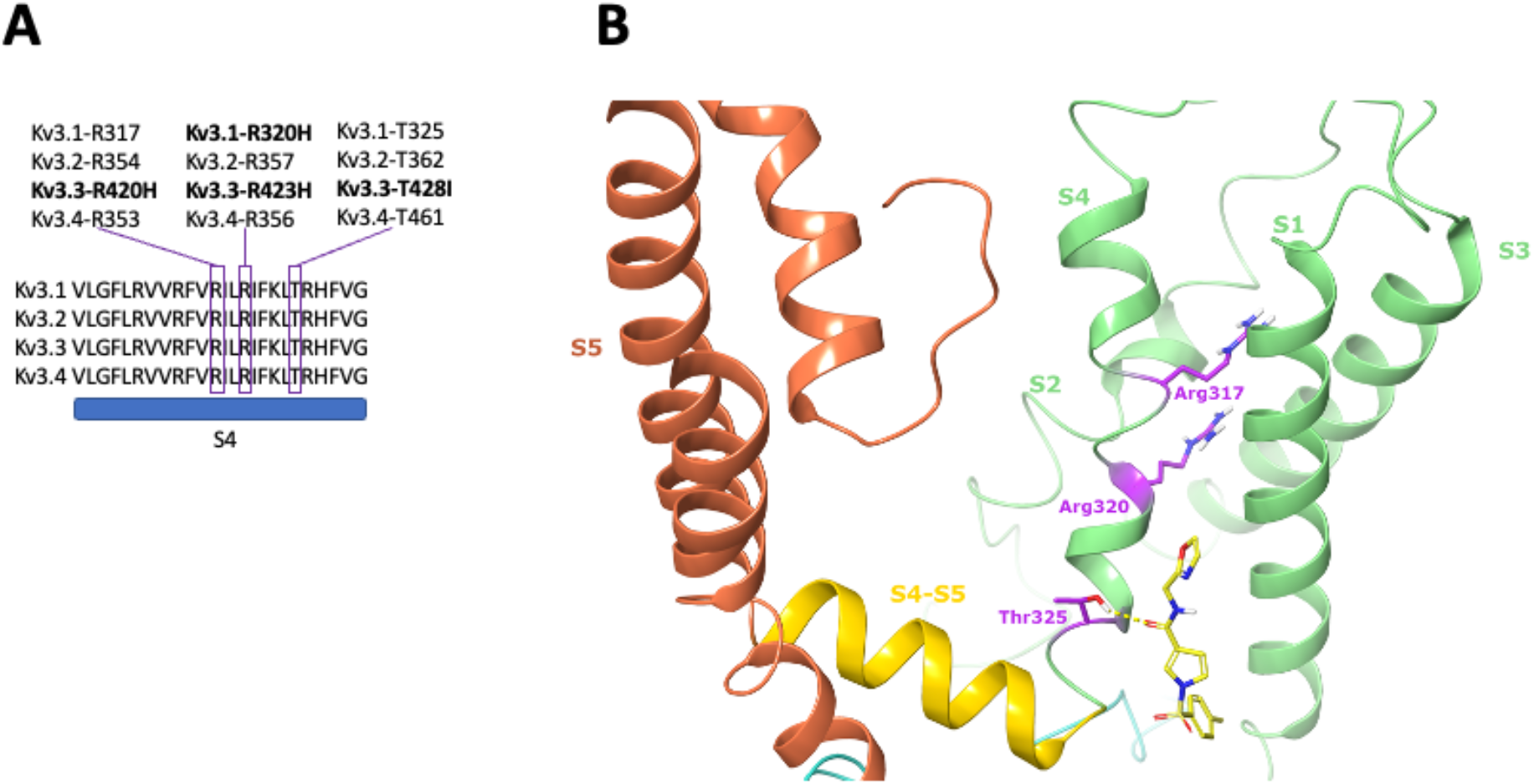
Region of disease related mutations in S4 in the VSD that were reported in the Kv3 class. **A**. Sequence alignment of human Kv3.1, Kv3.2 and Kv3.3, showing a high degree of conservation in the sequence of S4 in the VSD. Reported disease related mutations occurring in a class member are shown in bold. The corresponding residue in another class member is indicated **B**. Representation of the disease relevant Arg317, Arg320, and Thr325 residues (purple) in S4 within the VSD (light green), in the vicinity of the Lu AG00563 ligand pocket.

## REFERENCES

Adams, P.D., Afonine, P. V., Bunkóczi, G., Chen, V.B., Davis, I.W., Echols, N., Headd, J.J., Hung, L.W., Kapral, G.J., Grosse-Kunstleve, R.W., et al. (2010). PHENIX: A comprehensive Python-based system for macromolecular structure solution. Acta Crystallogr. Sect. D Biol. Crystallogr. 66, 213–221.

Andrade-Talavera, Y., Arroyo-García, L.E., Chen, G., Johansson, J., and Fisahn, A. (2020). Modulation of Kv3.1/Kv3.2 promotes gamma oscillations by rescuing Aβ-induced desynchronization of fast-spiking interneuron firing in an AD mouse model in vitro. J. Physiol. 598, 3711–3725.

Biyani, N. et al. Focus: The interface between data collection and data processing in cryo-EM. J. Struct. Biol.198, 124–133 (2017)

Brown, M.R., El-Hassar, L., Zhang, Y., Alvaro, G., Large, C.H., and Kaczmarek, L.K. (2016). Physiological modulators of Kv3.1 channels adjust firing patterns of auditory brain stem neurons. J. Neurophysiol. 116, 106–121.

England, S.K., Uebele, V.N., Shear, H., Kodali, J., Bennett, P.B., and Tamkun, M.M. (1995). Characterization of a voltage-gated K+ channel β subunit expressed in human heart. Proc. Natl. Acad. Sci. U. S. A. 92, 6309–6313.

Gallego-Iradi, C., Bickford, J.S., Khare, S., Hall, A., Nick, J.A., Salmasinia, D., Wawrowsky, K., Bannykh, S., Huynh, D.P., Rincon-Limas, D.E., Pulst, S.M., Harry, S.N., Fernandez-Funez, P., and Waters, M. (2014). KCNC3R420H, a K+ Channel Mutation Causative in Spinocerebellar Ataxia 13 Displays Aberrant Intracellular Trafficking. Neurobiol Dis. 71, 270–279

K. Nakahira, G. Shi K. Rhodes J.T. (1996). Selective Interaction of Voltage-gated K+ Channel beta-subunits with alpha-subunits. 271, 7084–7089.

Kaczmarek, L.K., and Zhang, Y. (2017). Kv3 channels: Enablers of rapid firing, neurotransmitter release, and neuronal endurance. Physiol. Rev.

Lewis, A., McCrossan, Z.A., and Abbott, G.W. (2004). MinK, MiRP1, and MiRP2 Diversify Kv3.1 and Kv3.2 Potassium Channel Gating. J. Biol. Chem. 279, 7884–7892.

Li, T., Wu, K., Yue, Z., Wang, Y., Zhang, F., and Shen, H. (2021a). Structural Basis for the Modulation of Human KCNQ4 by Small-Molecule Drugs. Mol. Cell.

Li, X., Zhang, Q., Guo, P., Fu, J., Mei, L., Lv, D., Wang, J., Lai, D., Ye, S., Yang, H., et al. (2021b). Molecular basis for ligand activation of the human KCNQ2 channel. Cell Res.

Liebschner, D. et al. Macromolecular structure determination using X-rays, neutrons and electrons: Recent developments in Phenix. Acta Crystallogr. Sect. D Struct. Biol. 75, 861–877 (2019).

Long, S.B., Campbell, E.B., and MacKinnon, R. (2005a). Crystal structure of a mammalian voltage-dependent Shaker family K + channel. Science (80-.). 309, 897–903.

Long, S.B., Campbell, E.B., and MacKinnon, R. (2005b). Voltage sensor of Kv1.2: Structural basis of electromechanical coupling. Science (80-.). 309, 903–908.

Long, S.B., Tao, X., Campbell, E.B., and MacKinnon, R. (2007). Atomic structure of a voltage-dependent K+ channel in a lipid membrane-like environment. Nature 450, 376–382.

Luneau, C.J., Williams, J.B., Marshall, J., Levitan, E.S., Oliva, C., Smith, J.S., Antanavage, J., Folander, K., Stein, R.B., Swanson, R., et al. (1991). Alternative splicing contributes to K+ channel diversity in the mammalian central nervous system. Proc. Natl. Acad. Sci. U. S. A. 88, 3932–3936.

Macica, C.M., Von Hehn, C.A.A., Wang, L.Y., Ho, C.S., Yokoyama, S., Joho, R.H., and Kaczmarek, L.K. (2003). Modulation of the Kv3.1b potassium channel isoform adjusts the fidelity of the firing pattern of auditory neurons. J. Neurosci. 23, 1133–1141.

Majumder, K., De Biasi, M., Wang, Z., and Wible, B.A. (1995). Molecular cloning and functional expression of a novel potassium channel β-subunit from human atrium. FEBS Lett.

Matthies, D., Bae, C., Toombes, G.E.S., Fox, T., Bartesaghi, A., Subramaniam, S., and Swartz, K.J. (2018). Single-particle cryo-EM structure of a voltage-activated potassium channel in lipid nanodiscs. Elife 7, 1–18.

McCrossan, Z.A., Lewis, A., Panaghie, G., Jordan, P.N., Christini, D.J., Lerner, D.J., and Abbott, G.W. (2003). MinK-related peptide 2 modulates Kv2.1 and Kv3.1 potassium channels in mammalian brain. J. Neurosci. 23, 8077–8091.

Morales, M.J., Castellino, R.C., Crews, A.L., Rasmusson, R.L., and Strauss, H.C. (1995). A novel β subunit increases rate of inactivation of specific voltage-gated potassium channel α subunits. J. Biol. Chem.

Muona, M., Berkovic, S.F., Dibbens, L.M., Oliver, K.L., Maljevic, S., Bayly, M.A., Joensuu, T., Canafoglia, L., Franceschetti, S., Michelucci, R., et al. (2015). A recurrent de novo mutation in KCNC1 causes progressive myoclonus epilepsy. Nat. Genet.

Németh, A.H., Kwasniewska, A.C., Lise, S., Parolin Schnekenberg, R., Becker, E.B.E., Bera, K.D., Shanks, M.E., Gregory, L., Buck, D., Zameel Cader, M., et al. (2013). Next generation sequencing for molecular diagnosis of neurological disorders using ataxias as a model. Brain 136, 3106–3118.

Oliver, K.L., Franceschetti, S., Milligan, C.J., Muona, M., Mandelstam, S.A., Canafoglia, L., Boguszewska-Chachulska, A.M., Korczyn, A.D., Bisulli, F., Di Bonaventura, C., et al. (2017). Myoclonus epilepsy and ataxia due to KCNC1 mutation: Analysis of 20 cases and K+ channel properties. Ann. Neurol.

Park, J., Koko, M., Hedrich, U.B.S., Hermann, A., Cremer, K., Haberlandt, E., Grimmel, M., Alhaddad, B., Beck-Woedl, S., Harrer, M., et al. (2019). KCNC1-related disorders: new de novo variants expand the phenotypic spectrum. Ann. Clin. Transl. Neurol. 6, 1319–1326.

Punjani, A., Rubinstein, J. L., Fleet, D. J. & Brubaker, M. A. CryoSPARC: Algorithms for rapid unsupervised cryo-EM structure determination. Nat. Methods 14, 290–296 (2017).

Rettig, J., Heinemann, S.H., Wunder, F., Lorra, C., Parcej, D.N., Oliver Dolly, J., and Pongs, O. (1994). Inactivation properties of voltage-gated K+ channels altered by presence of β-subunit. Nature.

Robertson, M.J., van Zundert, G.C.P., Borrelli, K., and Skiniotis, G. (2020). GemSpot: A Pipeline for Robust Modeling of Ligands into Cryo-EM Maps. Structure 28, 707-716.e3.

Rodriguez-Menchaca, A.A., Adney, S.K., Tang, Q.Y., Meng, X.Y., Rosenhouse-Dantsker, A., Cui, M., and Logothetis, D.E. (2012). PIP2 controls voltage-sensor movement and pore opening of Kv channels through the S4–S5 linker. Proc. Natl. Acad. Sci. U. S. A. 109, E2399–E2408.

Roos, K., Wu, C., Damm, W., Reboul, M., Stevenson, J.M., Lu, C., Dahlgren, M.K., Mondal, S., Chen, W., Wang, L., et al. (2019). OPLS3e: Extending Force Field Coverage for Drug-Like Small Molecules. J. Chem. Theory Comput. 15, 1863–1874.

Rohou, A. & Grigorieff, N. CTFFIND4: Fast and accurate defocus estimation from electron micrographs. J. Struct. Biol. 192, 216–221 (2015).

Rosato-Siri, M.D., Zambello, E., Mutinelli, C., Garbati, N., Benedetti, R., Aldegheri, L., Graziani, F., Virginio, C., Alvaro, G., and Large, C.H. (2015). A novel modulator of Kv3 potassium channels regulates the firing of parvalbumin-positive cortical interneuronss. J. Pharmacol. Exp. Ther.

Rudy, B., and McBain, C.J. (2001). Kv3 channels: Voltage-gated K+ channels designed for high-frequency repetitive firing. Trends Neurosci.

Sams, A.G., Rasmussen, L.K., Yu, W., and Fleming, P.R. Arylsulfonylpyrolecarboxamide derivatives as KV3 potassium channel activators. WO 2020/089262 A1. 2020 May 7

Scannevin, R.H., Wang, K.W., Jow, F., Megules, J., Kopsco, D.C., Edris, W., Carroll, K.C., Lü, Q., Xu, W., Xu, Z., et al. (2004). Two N-Terminal Domains of Kv4 K+ Channels Regulate Binding to and Modulation by KChIP1. Neuron 41, 587–598.

Scheres, S. H. W. & Chen, S. Prevention of overfitting in cryo-EM structure determination. Nat. Methods 9, 853–854 (2012).

Sun, J., and MacKinnon, R. (2017). Cryo-EM Structure of a KCNQ1/CaM Complex Reveals Insights into Congenital Long QT Syndrome. Cell 169, 1042-1050.e9.

Sun, J., and MacKinnon, R. (2020). Structural Basis of Human KCNQ1 Modulation and Gating. Cell 180, 340-347.e9.

Uhlhaas, P.J., and Singer, W. (2006). Neural Synchrony in Brain Disorders: Relevance for Cognitive Dysfunctions and Pathophysiology. Neuron 52, 155–168.

Wang, L.Y., Gan, L., Forsythe, I.D., and Kaczmarek, L.K. (1998). Contribution of the Kv3.1 potassium channel to high frequency firing in mouse auditory neurones. J. Physiol. 509, 183–194.

Zheng, S. Q. et al. MotionCor2: Anisotropic correction of beam-induced motion for improved cryo-electron microscopy. Nat. Methods 14, 331–332 (2017).

Zhu, K. et al. Docking covalent inhibitors: A parameter free approach to pose prediction and scoring. J. Chem. Inf. Model. 54, 1932–1940 (2014).

